# Overexpression of Interleukin-8 and Interleukin-13 as potent immune markers associated with survival and dietary habits in esophageal squamous cell carcinoma

**DOI:** 10.1101/2022.10.17.512528

**Authors:** Jayasree Talukdar, Abdul Malik, Kangkana Kataki, Bikash Narayan Choudhury, Munindra Narayan Baruah, Mallika Bhattacharyya, Musaed Alkholief, Aws Alshamsan, Manash Pratim Sarma, Minakshi Bhattacharjee, Mrinmoy Basak, Manash Pratim Kashyap, Sahana Bhattacharjee, Eyashin Ali, Chenole Keppen, Mohammad Ghaznavi Idris, Subhash Medhi

**Affiliations:** Department of Bioengineering and Technology, Gauhati University, Gawahati, Assam, India; Department of Pharmaceutics, College of Pharmacy, King Saud University, Riyadh, Saudi Arabia; Department of Gastroentrology, Gauhati Medical College Hospital, Guwahati, Assam, India; Department of Head and Neck Oncology, North East Cancer Hospital and Research Institute, Jorabat, Assam, India; Department of Biotechnology, Faculty of science, Assam down town University, Guwahati, Assam, India; Faculty of pharmaceutical Sciences, Assam down town University, Guwahati, Assam, India; Department of Statistics, Faculty of science, Assam down town University, Guwahati, Assam, India; Department of Statistics, Gauhati University, Gawahati, Assam, India

**Keywords:** Interleukin-8, Interleukin-12, Interleukin-13, Esophageal squamous cell carcinoma, survival analysis, dietary habits

## Abstract

**Background:** Interleukin-8 (IL8), Interleukin-12 (IL12) and Interleukin-13 (IL13) are cytokines that play regulatory role in cancer pathogenesis. We analysed their expression profile to evaluate as molecular biomarkers of esophageal squamous cell carcinoma (ESCC) and their association with different parameters.

**Methods:** Expression analysis of IL8, IL12 and IL13 were performed by Real time qPCR in blood and tumor tissue of 120 ESCC patients. The expression profiles were associated with different clinicopathological and dietary factors. Survival and hazard analysis were also performed.

**Results:** When compared to normal controls, IL8 expression showed upregulation in 83% tissue samples (p=0.000) and 62% blood samples (p=0.388), IL12 expression showed upregulation in 62% tissue samples (p=0.435) and 57% blood samples (p=0.222) and IL13 expression showed upregulation in 83% tissue samples (p=0.001) and 68% blood samples (p=0.312). Significant positive correlation (p<0.05) was observed between tissue and blood level expression of IL8, IL12 and IL13. Different clinicopathological factors and dietary habits showed significant association (p<0.05) with IL8, IL12 and IL13 expression.. Statistically significant positive correlation were observed for IL8 and IL13 expression in tissue as well as IL13 and IL12 expression in both tissue and blood. Also significant negative correlation of IL8 and IL12 expression in blood and tissue were also observed. Tumor stage, node stage, metastasis, consumption of betel nut, tobacco, alcohol, hot food, smoked food, spices, IL8 expression in blood, IL13 expression in tissue and IL12 expression in blood and tissue showed significant association (p<0.05) with survival of ESCC patients.

**Conclusions:** Altered expression of IL8, IL12 and IL13 may be associated with ESCC progression. Overexpression of IL8 and IL13 in tissue samples may be potential biomarkers for ECSS screening. Additionally, results from both survival and hazard analysis data indicate the effects of various parameters on the survival and mortality rate of ESCC patients.

## Introduction

Esophageal cancer represents the seventh most common cancer worldwide and it is the sixth most common reason of cancer-related death worldwide with a survival rate of only 15-20 % in five years [1, 2]. Esophageal squamous cell carcinoma (ESCC) is reported as the most prevalent histological form of esophageal cancer in the world [3]. Patients with ESCC often present at an advanced stage when diagnosed because there are ineffective early detection tools, which has a negative impact on the patients’ prognosis. Detection of molecular biomarkers might open up new, efficient means for tumour diagnosis, screening, monitoring, and prognosis [4, 5].

Tumor microenvironment consists of many different types of cells, which produces different kind of cytokines that can either enhance or inhibit cancer cell proliferation. These all interact with one another to play their role in tumor pathogenesis [6, 7]. Interleukin-8 (IL8) is a pro-inflammatory cytokine produced by different cell types in response to tissue infection, inflammation or injury [8-10]. It is also associated with the development of different types of cancer like lung cancer, prostate cancer, breast cancer, colorectal cancer, etc. [8, 11]. It plays a dual potential role in tumor microenvironment by directly promoting tumor survival and indirectly facilitating tumor progression by affecting components of tumor microenvironment, which include epithelial-to-mesenchymal transition, pro-angiogenesis process, tumor cell proliferation and inhibition of anti-tumor immunity [8]. Interleukin-12 (IL12), an essential pro-inflammatory heterodimeric cytokine, is primarily produced by antigen presenting cells in response to infection. It plays an important role in connecting adaptive and innate arm of immune systems and stimulates the activity of natural killer cell and T cell and induces production of interferon gamma [12, 13]. It is a potent agent in enhancing antitumor immune responses and plays important roles in the regulation of cellular immunity [14]. It has been considered as an essential immunotherapeutic agent for combinatorial cancer treatments [13]. Interleukin 13 (IL-13) is an immunoregulatory cytokine, synthesized primarily by activated T-helper 2 cells, but also by B cells, natural killer, dendritic cells, mastocytes, basophils, etc. [15, 16]. IL13 and its receptors play an essential role in the proliferation of cancer cells and other biological behaviours like invasion, migration etc. and enhance the malignant phenotype. In many human cancers, the presence of IL13 and its receptors are reported to have association with chemosensitivity, apotosis and cancer prognosis [17-19]. In this study we have analysed the expression of cytokine IL8, IL12 and IL13 between blood and tissue samples and also among male and female patients of ESCC. We further evaluated the association between the expression profile of these cytokines and different clinicopathological and dietary factors in ESCC patients.

The survival rate of esophageal cancer is quite low because most cases of this cancer are detected at a late stage. Several factors affect the survival of esophageal cancer patients and these factors may vary in esophageal cancer patients from different populations. Therefore, research that identifies esophageal cancer risk factors is necessary to enhance patient prognosis and survival rates [20]. In this study, survival and hazard analysis were performed to check the impact of different risk factors on survival or mortality rate of ESCC patients.

## Materials and Methods

### Sample collection

Blood samples were collected using standard venipuncture and tumor tissues as well as adjacent normal tissues were collected by biopsy from 120 ESCC patients with written informed consent. The diagnosis of ESCC was confirmed by upper gastrointestinal endoscopy (UGI endoscopy) and by pathological analysis of tumor biopsies. An equal number of healthy individuals (age and sex matched) were also enrolled in the study. The study was approved by the Institutional Ethical Committee of the Gauhati Medical College and Hospital, Guwahati (MC/217/2016/Pt-I/20; dated 22 December 2016) and North East Cancer Hospital, Jorabat (IEC/2018/06/NP/12; dated 27 August 2018) and all the procedures were in accordance with the Helsinki Declarations and with the ethical standards of the Institutional Ethical Committee. Histopathology grade, stages of tumor, node and metastasis were categorized according to AJCC (American Joint Committee on Cancer) manual of cancer staging. Dysphagia grade was determined using the modified O’Rourke grading system. Amount of tea consumption was categorized into low (who consumed1-2 times/day), medium (who consumed 3-4times/day) and high amount (who consumed 5 or more times/day). Amount of khar consumption was categorized into no (who don’t consume), low (1-2 times/month), medium (3-4 times/month) and high (5 or more times/month) amount.

### mRNA expression profile analysis by Real Time PCR

RNA isolation was done manually using the Trizol reagent from homogenized tissue and blood samples according to the manufacturer’s instruction. Reverse transcription was done to prepare complementary DNA (cDNA) using iScript™ Reverse Transcription Reagents (Bio-Rad Laboratories, Inc.). Real time qPCR was performed using SYBR green master mix in a Rotor-Gene Q (Qiagen) for analysis of mRNA expression. Human housekeeping gene β-actin was considered as standard reference for normalization. The level of expression of the targeted genes was calculated using the formula Comparative Ct (2^-ΔΔCt^) method. If the calculated level of expression was <1, the targeted gene was downregulated and if the level of expression was >1, the targeted gene was upregulated. The primer used for IL8, IL13, IL12 and β-actin genes were: Forward (F): 5′-TCTGTCTGGACCCCAAGGAA-3′, Reverse (R): 5-GCAACCCTACAACAGACCCA-3′; F: 5′-GCACAGACCAAGGCAAATG-3′, R: 5′-GCAGAATGAGTGCTGTGGA-3′; F: 5′-TGATGAAGAAGCTGCTGGT-3′, R: 5′-GTCAGAGGGGACAACA-3′ and F: 5′-AGATGTGGATCAGCAAGCAG-3′, R: 5′-GCGCAAGTTAGGTTTTGTCA-3′ respectively.

### Statistical analysis

The Statistical Package for the Social Sciences (SPSS) version 18.0 was used for all statistical analysis. All the data were expressed as mean ± standard deviation and two tailed tests were taken into consideration. All the tests were considered statistically significant with a p-value < 0.05. Student’s paired t test was used to compare mRNA expression levels in cases and controls. The parametric independent sample’s t test and one way ANOVA or the non-parametric Mann-Whitney U test and Kruskal Wallis H test were performed to study the association with different parameters. Kaplan–Meier method of survival analysis was done using the log-rank test and univariate model of Cox’s regression was used to detect hazard outcomes for different risk factors.

## Results

### IL8, IL12 and IL13 expression in blood and tissue samples of ESCC patients

IL8 expression was upregulated in 83% ESCC cases with a mean fold change of 2.63±1.06 and downregulated (0.40±0.24) in 17% ESCC cases in tissue samples (p=0.000). In blood samples, IL8 expression was upregulated (2.37±1.19) in 62% cases and downregulated (0.28±0.22) in 38% cases (p=0.388). While analyzing IL12 expression, 62% cases showed upregulation (1.77±0.75) and 38% cases showed downregulation (0.46±0.31) in tissue samples (p=0.435). In blood samples, 57% cases showed upregulation (2.10±0.75) and 43% cases showed downregulation (0.42±0.30) of IL12 (p=0.227). Expression of IL13 was upregulated (2.27±0.91) in 83% cases and downregulated (0.46±0.26) in 17% cases in tissue samples (p=0.001). While analyzing blood samples, 68% cases showed upregulation (2.30±1.08) and 32% cases showed downregulation (0.49±0.30) of IL13 (p=0.312). Tissue and blood level expression of IL8, IL12 and IL13 in ESCC patients are represented in Table 1. Box plot representation of IL8, IL12 and IL13 expression in ESCC patients are represented in Fig 1 and box plot representation of IL8, IL12 and IL13 expression in ESCC patients compared to normal control are represented in Fig 2.

**Table 1.**
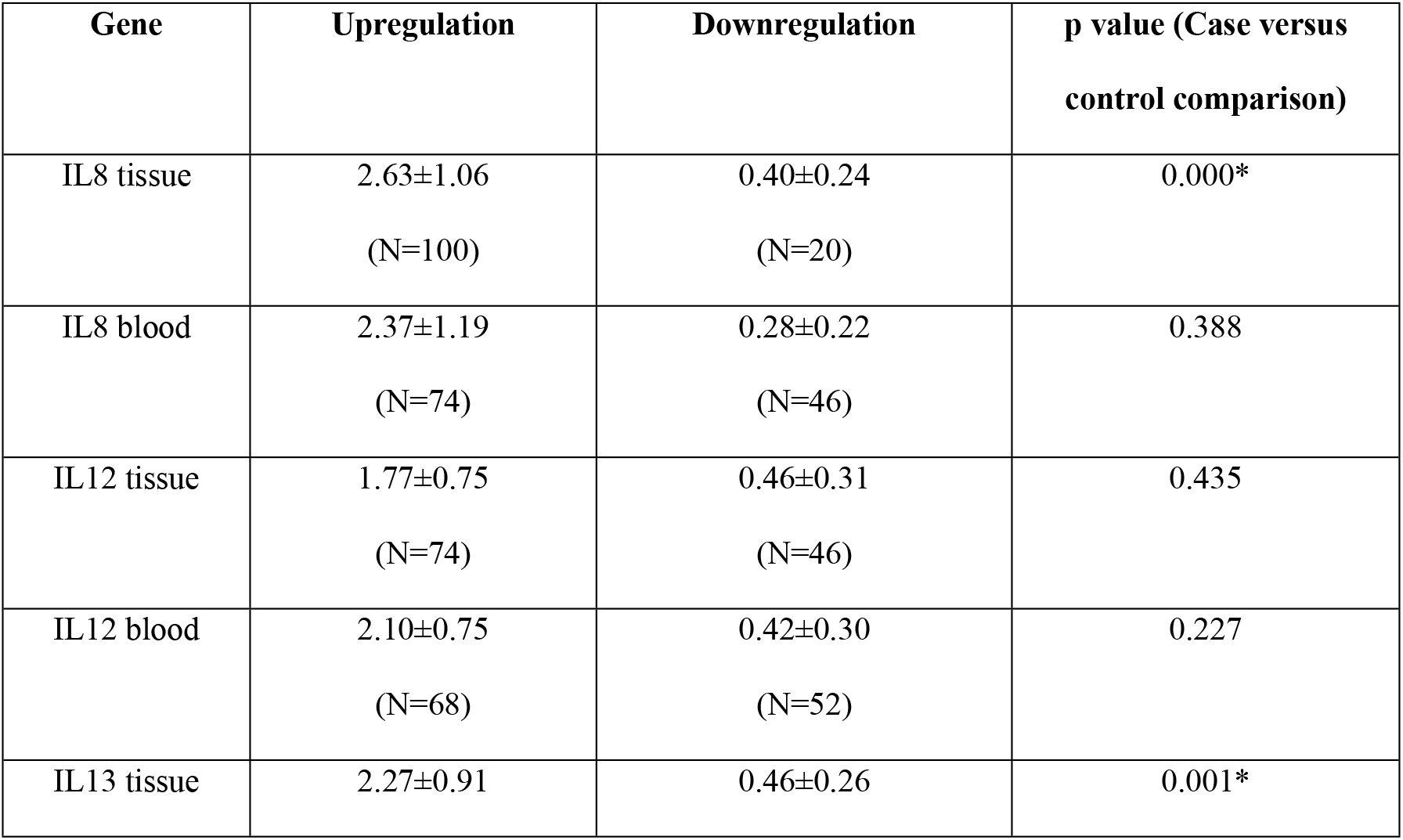
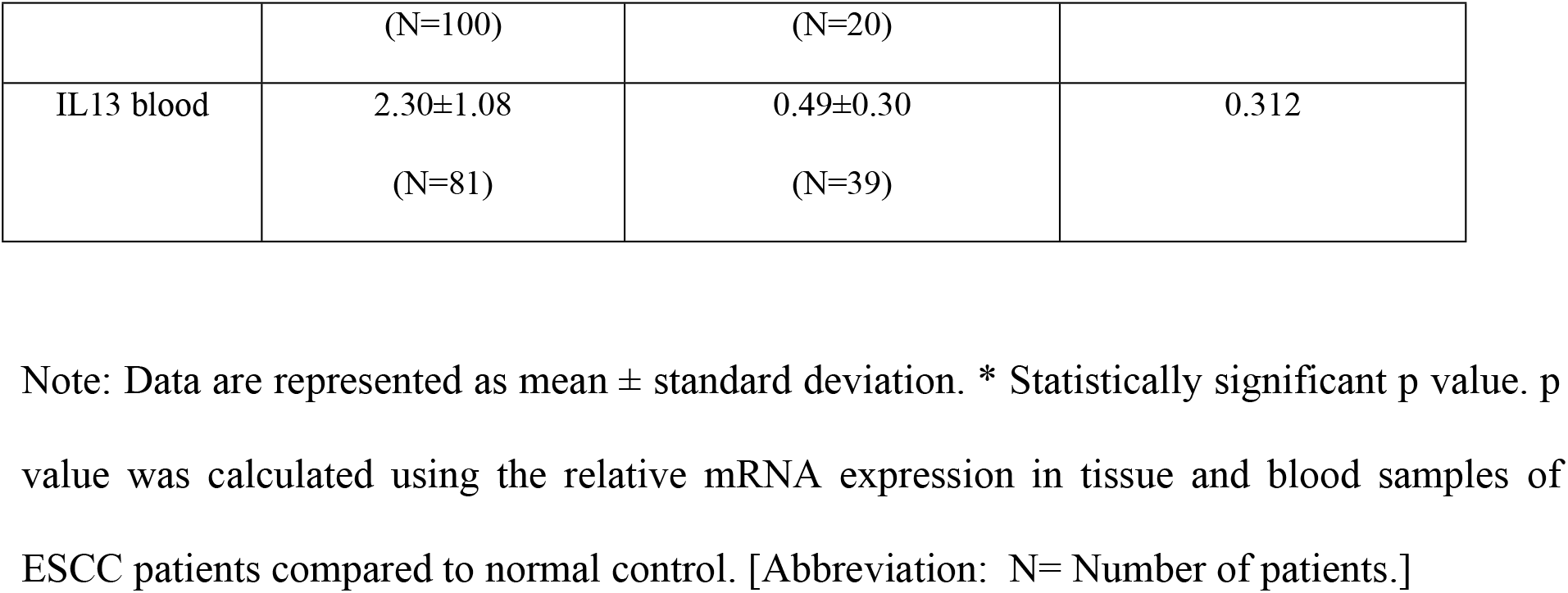
Expression profile of IL8, IL13 and IL12 gene in both blood and tissue of ESCC patients.

**Fig 1.**
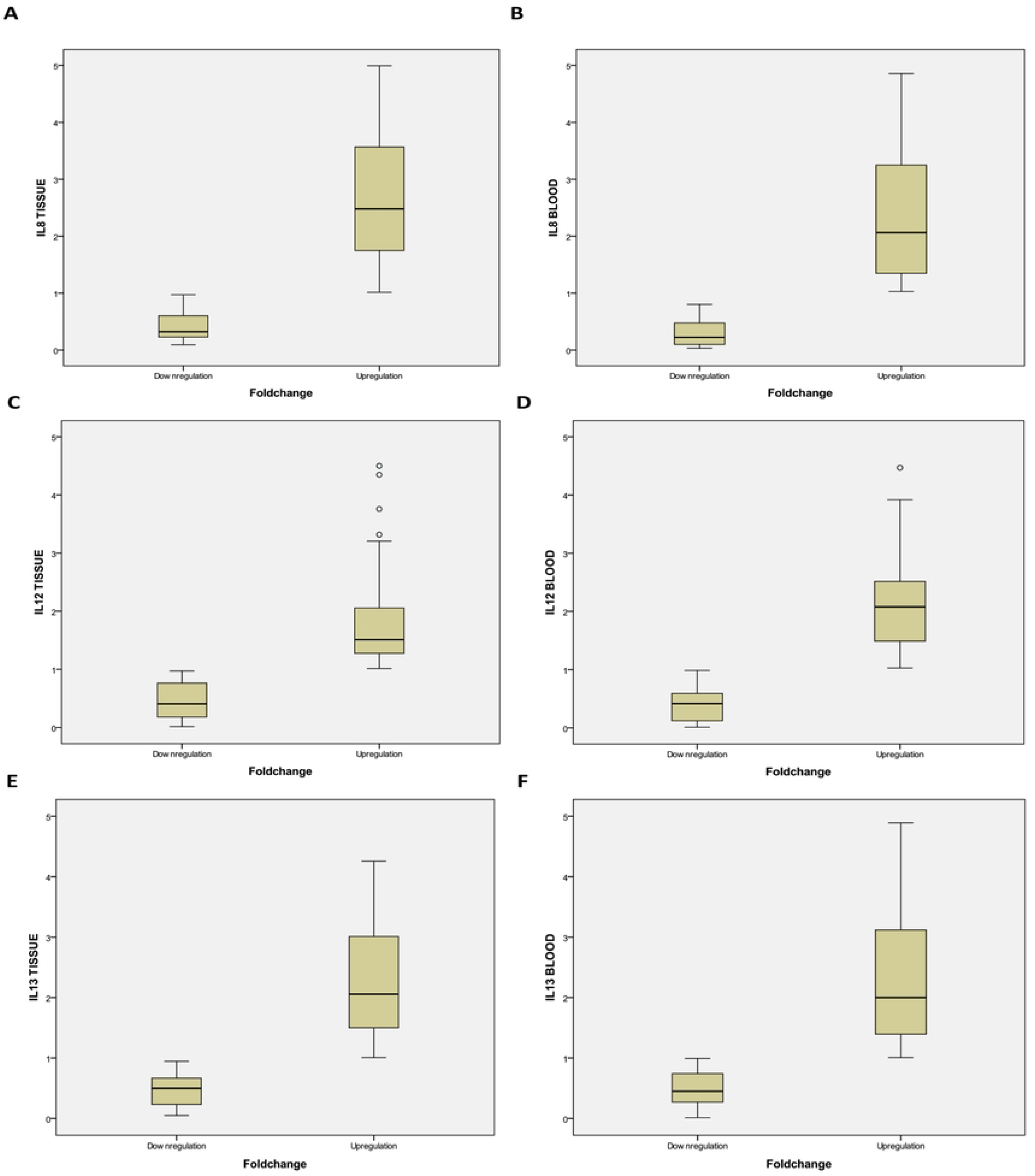
Box plot representation of IL8, IL12 and IL13 expression in tissue and blood level of ESCC patients. (A) IL8 expression in tissue; (B) IL8 expression in blood; (C) IL13 expression in tissue; (D) IL13 expression in blood; (E) IL12 expression in tissue; (F) IL12 expression in blood. [Note: Box plot explanation: Dark horizontal bar within box–median; upper horizontal line of box-75th percentile; lower horizontal line of box-25th percentile; whiskers and dots-range in the box plot.]

**Fig 2.**
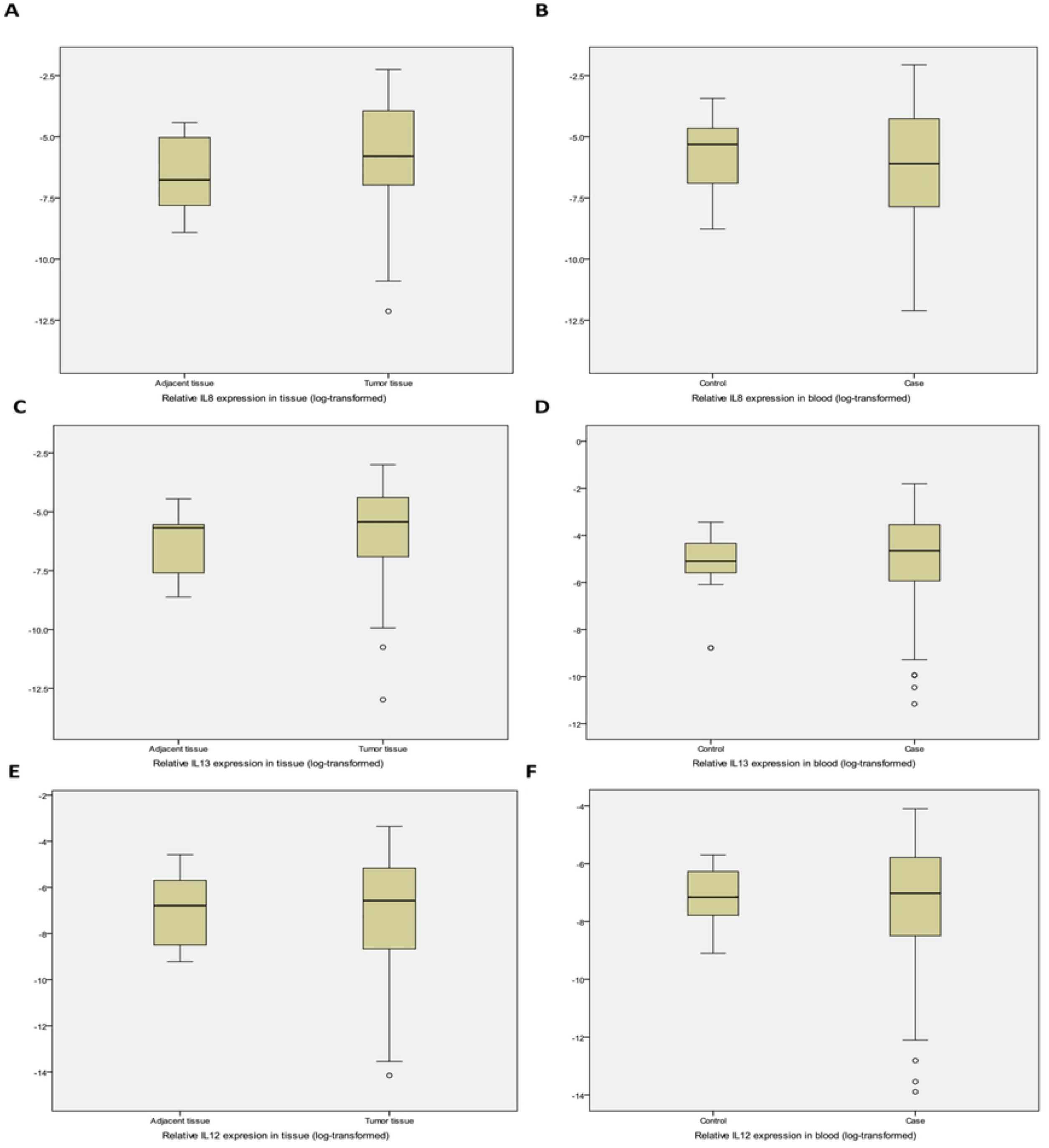
Box plot representation of IL8, IL12 and IL13 expression in tissue and blood samples of ESCC patients compared to normal control. (A) Relative IL8 expression in tumor tissues and adjacent normal tissues; (B) Relative IL8 expression in blood samples and control samples; (C) Relative IL13 expression in tumor tissues and adjacent normal tissues; (D) Relative IL13 expression in blood samples and control samples; (E) Relative IL12 expression in tumor tissues and adjacent normal tissues; (F) Relative IL12 expression in blood samples and control samples. [Note: Box plot explanation: Dark horizontal bar within box–median; upper horizontal line of box-75th percentile; lower horizontal line of box-25th percentile; whiskers and dots-range in the box plot.]

### Association of IL8, IL12 and IL13 expression with different clinicopathological and dietary factors in ESCC

While targeting the IL8 expression, significant associations (p<0.05) were observed in age group, dysphagia grade, metastasis, consumption of alcohol, betel nut, spices, meat and hot food in tissue samples of male cases (S1 Fig), while age group, dysphagia grade, tumor stage, consumption of betel nut, hot food, smoked food, spices and amount of tea consumed showed significant differences in female cases (S2 Fig). For blood samples, male patients showed significant association with age group, dysphagia grade, consumption of tobacco, betel nut, hot food, spices and type of tea consumed (S3 Fig), whereas female patients showed an association with age group, dysphagia grade, tumor stage, consumption of betel nut, spices and hot food (S4 Fig). For IL12 expression in tissue samples, male cases showed significant association with age group, habit of smoking, consumption of smoked food, fast food, tobacco, spices, hot food and types of tea consumed (S5 Fig) while female cases showed association with age group, consumption of betel nut, hot food, smoked food, and spices (S6 Fig). While considering blood samples, IL12 expression showed association with age group, habit of smoking, consumption of smoked food, fast food, hot food and spices in male cases (S7 Fig) and female cases were associated with age group, tumor stage, metastasis, consumption of smoked food and spices and amount of tea consumed (S8 Fig). For IL13 expression, significant associations were observed in histopathology grade, dysphagia grade, location of tumor, consumption of tobacco, hot food and smoked food in tissue samples of male patients (S9 Fig). Among females, significant difference was found in histopathology grade, tumor stage, habit of smoking, consumption of tobacco, fish, meat and smoked food in their tissue samples (S10 Fig). While checking blood samples, male cases showed significant association with histopathology grade, consumption of tobacco, amount of tea consumed and types of tea consumed (S11 Fig) while female cases were found to associate with histopathology grade, consumption of tobacco and khar and differences in the amount of khar consumption (S12 Fig). Association study of IL8, IL12 and IL13 expression with 24 different parameters are listed in Table 2 (for males) and Table 3 (for females).

**Table 2.** Tabulation of association of IL8, IL12 and IL13 expression with different clinicopathological parameters and lifestyle factors in male ESCC patients.

| Clinicopathological parameters and lifestyle factors | No of cases (%) | IL8 tissue |  | IL8 blood |  | IL13 tissue |  | IL13 blood |  | IL12tissue |  | IL12 blood |  |
| --- | --- | --- | --- | --- | --- | --- | --- | --- | --- | --- | --- | --- | --- |
|  |  | mRNA expression | P value | mRNA expression | P value | mRNA expression | P value | mRNA expression | P value | mRNA expression | P value | mRNA expression | P value |
| Age group: |  |  | 0.004* |  | 0.001* |  | 0.175 |  | 0.297 |  | 0.003* |  | 0.012* |
| <50years | 29(41%) | 1.70±1.00 |  | 1.06±1.27 |  | 1.74±0.90 |  | 1.69±1.26 |  | 1.75±1.09 |  | 1.69±1.01 |  |
| >50years | 41(59%) | 2.58±1.35 |  | 2.02±1.36 |  | 2.10±1.20 |  | 2.01±1.31 |  | 1.03±0.73 |  | 1.12±1.08 |  |
| Gender: |  |  | 0.648 |  | 0.539 |  | 0.843 |  | 0.106 |  | 0.571 |  | 0.707 |
| Male | 70(58%) | 2.22±1.29 |  | 1.62±1.40 |  | 1.95±1.09 |  | 1.88±1.29 |  | 1.33±0.96 |  | 1.33±0.96 |  |
| Female | 50(42%) | 2.33±1.28 |  | 1.50±1.37 |  | 1.99±1.06 |  | 1.48±1.14 |  | 1.19±0.78 |  | 1.39±0.96 |  |
| Location of tumor: |  |  | 0.655 |  | 0.070 |  | 0.036* |  | 0.273 |  | 0.811 |  | 0.918 |
| Upper esophagus | 19(97%) | 2.14±1.53 |  | 1.46±1.32 |  | 2.48±0.97 |  | 1.69±1.27 |  | 1.20±0.83 |  | 1.31±1.18 |  |
| Middle esophagus | 24(34%) | 2.08±1.05 |  | 1.25±1.25 |  | 1.85±1.21 |  | 1.72±1.42 |  | 1.37±1.00 |  | 1.37±1.11 |  |
| Lower esophagus | 27(39%) | 2.40±1.33 |  | 2.06±1.51 |  | 1.66±0.95 |  | 2.15±1.18 |  | 1.38±1.02 |  | 1.37±1.03 |  |
| Histopathology grade: |  |  | 0.463 |  | 0.757 |  | 0.003* |  | 0.007* |  | 0.835 |  | 0.771 |
| Grade 1:well differentiated ESCC | 24(34%) | 1.96±0.96 |  | 1.68±1.49 |  | 1.38±0.80 |  | 1.41±1.15 |  | 1.26±0.89 |  | 1.43±1.20 |  |
| Grade 2: moderately differentiated ESCC | 35(50%) | 2.33±1.40 |  | 1.47±1.27 |  | 2.14±1.10 |  | 1.88±1.27 |  | 1.35±1.05 |  | 1.27±1.03 |  |
| Grade 3:poorly differentiated ESCC | 11(16%) | 2.44±1.55 |  | 1.96±1.65 |  | 2.59±1.13 |  | 2.89±1.13 |  | 1.40±0.87 |  | 1.47±1.05 |  |
| Dysphagia grade: |  |  | 0.008* |  | 0.012* |  | 0.026* |  | 0.633 |  | 0.214 |  | 0.543 |
| Grade 0: Asymptomatic | 02(03%) | 1.71±2.07 |  | 0.64±0.57 |  | 2.56±0.99 |  | 1.85±1.21 |  | 1.17±0.32 |  | 0.68±0.93 |  |
| Grade 1: Solids with some dysphagia | 23(33%) | 1.60±1.09 |  | 0.81±0.57 |  | 2.09±0.85 |  | 1.76±1.50 |  | 1.52±1.06 |  | 1.63±1.16 |  |
| Grade 2: Soft, pureed food only | 07(10%) | 2.01±1.21 |  | 1.22±1.10 |  | 2.72±1.33 |  | 2.06±1.42 |  | 1.76±0.99 |  | 1.35±0.96 |  |
| Grade 3:Liquids only | 31(44%) | 2.48±1.26 |  | 2.09±1.43 |  | 1.52±0.99 |  | 1.77±1.08 |  | 1.13±0.94 |  | 1.19±1.07 |  |
| Grade 4: No swallowing at all | 07(10%) | 3.41±1.04 |  | 2.86±2.00 |  | 2.45±1.44 |  | 2.60±1.42 |  | 1.16±0.58 |  | 1.37±1.11 |  |
| Tumor stage: |  |  | 0.818 |  | 0.955 |  | 0.552 |  | 0.311 |  | 0.126 |  | 0.789 |
| Stage1 | 03(04%) | 2.45±2.25 |  | 1.38±0.21 |  | 1.61±0.33 |  | 0.69±0.55 |  | 2.36±0.51 |  | 1.79±1.22 |  |
| Stage2 | 13(19%) | 2.17±1.20 |  | 1.90±1.80 |  | 2.32±0.92 |  | 2.00±1.27 |  | 1.15±0.74 |  | 1.47±1.07 |  |
| Stage3 | 35(50%) | 2.10±1.41 |  | 1.53±1.18 |  | 1.83±1.15 |  | 2.04±1.37 |  | 1.18±0.83 |  | 1.37±1.19 |  |
| Stage4 | 19(27%) | 2.44±0.99 |  | 1.63±1.62 |  | 1.97±1.16 |  | 1.69±1.19 |  | 1.54±1.24 |  | 1.18±0.91 |  |
| Node stage: |  |  | 0.908 |  | 0.652 |  | 0.650 |  | 0.542 |  | 0.958 |  | 0.314 |
| Stage 0 | 21(30%) | 2.13±1.40 |  | 1.75±1.62 |  | 1.77±0.85 |  | 1.65±1.01 |  | 1.29±0.81 |  | 1.55±1.33 |  |
| Stage1 | 25(36%) | 2.24±1.18 |  | 1.44±1.19 |  | 2.01±1.12 |  | 1.81±1.53 |  | 1.30±0.91 |  | 1.39±0.96 |  |
| Stage2 | 19(27%) | 2.36±1.36 |  | 1.77±1.31 |  | 1.94±1.35 |  | 2.17±1.16 |  | 1.31±1.03 |  | 1.30±1.01 |  |
| Stage3 | 05(07%) | 1.94±1.39 |  | 1.35±1.97 |  | 2.45±0.78 |  | 2.09±1.66 |  | 1.67±1.64 |  | 0.53±0.48 |  |
| Metastasis: |  |  | 0.036* |  | 0.511 |  | 0.702 |  | 0.590 |  | 0.476 |  | 0.095 |
| Absent | 54(77%) | 2.04±1.27 |  | 1.52±1.33 |  | 1.92±1.00 |  | 1.84±1.29 |  | 1.31±0.84 |  | 1.49±1.11 |  |
| Present | 16(33%) | 2.81±1.22 |  | 1.95±1.61 |  | 2.04±1.38 |  | 2.01±1.32 |  | 1.37±1.32 |  | 0.91±0.86 |  |
| Smoking: |  |  | 0.891 |  | 0.562 |  | 0.573 |  | 0.988 |  | 0.033* |  | 0.016* |
| Smokers | 39(56%) | 2.24±1.34 |  | 1.73±1.52 |  | 2.01±1.04 |  | 1.91±1.42 |  | 1.12±0.81 |  | 1.12±1.11 |  |
| Nonsmokers | 31(44%) | 2.19±1.25 |  | 1.48±1.25 |  | 1.86±1.17 |  | 1.83±1.12 |  | 1.59±1.08 |  | 1.65±0.98 |  |
| Alcohol: |  |  | 0.035* |  | 0.573 |  | 0.327 |  | 0.318 |  | 0.764 |  | 0.213 |
| Alcoholic | 36(51%) | 2.53±1.35 |  | 1.73±1.51 |  | 2.07±1.12 |  | 2.03±1.31 |  | 1.30±0.98 |  | 1.51±1.10 |  |
| Nonalcoholic | 34(49%) | 1.88±1.14 |  | 1.50±1.29 |  | 1.82±1.06 |  | 1.72±1.27 |  | 1.36±0.95 |  | 1.19±1.06 |  |
| Betel nut: |  |  | 0.002* |  | 0.009* |  | 0.464 |  | 0.548 |  | 0.821 |  | 0.178 |
| Chewers | 45(64%) | 2.56±1.25 |  | 1.98±1.54 |  | 2.02±1.08 |  | 1.81±1.29 |  | 1.38±1.07 |  | 1.47±1.10 |  |
| Nonchewers | 25(36%) | 1.60±1.14 |  | 0.97±0.79 |  | 1.82±1.13 |  | 1.32±0.82 |  | 1.23±0.73 |  | 1.14±1.03 |  |
| Tobacco: |  |  | 0.674 |  | 0.013* |  | 0.006* |  | 0.022* |  | 0.032* |  | 0.710 |
| Chewers | 46(66%) | 2.26±1.37 |  | 1.92±1.48 |  | 2.20±1.10 |  | 2.17±1.40 |  | 1.21±0.99 |  | 1.37±1.06 |  |
| Nonchewers | 24(34%) | 2.13±1.14 |  | 1.05±1.03 |  | 1.46±0.91 |  | 1.32±0.82 |  | 1.56±0.87 |  | 1.33±1.15 |  |
| Fish: |  |  | 0.740 |  | 0.456 |  | 0.522 |  | 0.069 |  | 0.288 |  | 0.648 |
| Consumers | 61(87%) | 2.20±1.22 |  | 1.66±1.41 |  | 1.98±1.10 |  | 1.98±1.29 |  | 1.27±0.93 |  | 1.39±1.11 |  |
| Nonconsumers | 09(13%) | 2.35±1.79 |  | 1.34±1.37 |  | 1.73±1.10 |  | 1.19±1.10 |  | 1.68±1.11 |  | 1.12±0.88 |  |
| Meat: |  |  | 0.008* |  | 0.417 |  | 0.635 |  | 0.056 |  | 0.108 |  | 0.283 |
| Consumers | 67(96%) | 2.30±1.25 |  | 1.66±1.42 |  | 1.96±1.09 |  | 1.81±1.28 |  | 1.27±0.92 |  | 1.33±1.10 |  |
| Nonconsumers | 03(04%) | 0.32±0.14 |  | 0.78±0.63 |  | 1.65±1.33 |  | 3.31±0.57 |  | 2.48±1.24 |  | 1.80±0.44 |  |
| Egg: |  |  | 0.392 |  | 0.223 |  | 0.307 |  | 0.337 |  | 0.128 |  | 0.328 |
| Consumers | 62(89%) | 2.27±1.30 |  | 1.69±1.41 |  | 1.90±1.06 |  | 1.92±1.26 |  | 1.25±0.88 |  | 1.32±1.09 |  |
| Nonconsumers | 08(11%) | 1.85±1.18 |  | 1.10±1.27 |  | 2.32±1.33 |  | 1.55±1.55 |  | 1.93±1.35 |  | 1.59±1.04 |  |
| Pickle: |  |  | 0.304 |  | 0.637 |  | 0.394 |  | 0.256 |  | 0.951 |  | 0.203 |
| Consumers | 53(76%) | 2.13±1.19 |  | 1.52±1.27 |  | 1.88±1.11 |  | 1.96±1.27 |  | 1.33±0.99 |  | 1.26±1.06 |  |
| Nonconsumers | 17(24%) | 2.50±1.56 |  | 1.94±1.76 |  | 2.15±1.04 |  | 1.62±1.37 |  | 1.31±0.86 |  | 1.64±1.12 |  |
| Spices: |  |  | 0.011* |  | 0.007* |  | 0.840 |  | 0.210 |  | 0.012* |  | 0.020* |
| Consumers | 53(76%) | 2.44±1.33 |  | 1.86±1.43 |  | 1.96±1.15 |  | 1.97±1.23 |  | 1.10±0.67 |  | 1.14±0.90 |  |
| Nonconsumers | 17(24%) | 1.53±0.86 |  | 0.87±1.02 |  | 1.90±0.92 |  | 1.60±1.48 |  | 2.02±1.35 |  | 2.02±1.35 |  |
| Fast food: |  |  | 0.500 |  | 0.381 |  | 0.518 |  | 0.788 |  | 0.015* |  | 0.007* |
| Consumers | 47(67%) | 2.29±1.30 |  | 1.67±1.35 |  | 1.89±1.13 |  | 1.91±1.33 |  | 1.12±0.83 |  | 1.07±0.86 |  |
| Nonconsumers | 23(33%) | 2.07±1.27 |  | 1.52±1.53 |  | 2.07±1.03 |  | 1.81±1.24 |  | 1.74±1.09 |  | 1.95±1.26 |  |
| Hot food: |  |  | 0.010* |  | 0.040* |  | 0.043* |  | 0.114 |  | 0.014* |  | 0.013* |
| Consumers | 43(61%) | 2.53±1.35 |  | 1.91±1.52 |  | 2.16±1.18 |  | 2.07±1.33 |  | 1.13±0.88 |  | 1.12±1.06 |  |
| Nonconsumers | 27(39%) | 1.72±1.02 |  | 1.15±1.05 |  | 1.62±0.86 |  | 1.57±1.18 |  | 1.64±1.01 |  | 1.73±1.03 |  |
| Type of tea: |  |  | 0.269 |  | 0.036 |  | 0.790 |  | 0.032* |  | 0.030* |  | 0.199 |
| Red tea | 18(26%) | 2.15±1.30 |  | 1.03±1.14 |  | 1.85±0.97 |  | 1.30±0.94 |  | 1.56±0.97 |  | 1.72±1.06 |  |
| Milk tea | 21(30%) | 1.89±1.24 |  | 1.98±1.46 |  | 2.08±0.95 |  | 2.42±1.38 |  | 1.03±1.08 |  | 1.23±0.98 |  |
| Both | 31(44%) | 2.48±1.30 |  | 1.72±1.43 |  | 1.92±1.26 |  | 1.85±1.29 |  | 1.39±0.83 |  | 1.23±1.15 |  |
| Amount of tea: |  |  | 0.545 |  | 0.269 |  | 0.832 |  | 0.005* |  | 0.114 |  | 0.192 |
| Low | 21(30%) | 2.29±1.44 |  | 1.22±1.21 |  | 1.88±1.02 |  | 1.14±0.95 |  | 1.56±0.86 |  | 1.78±1.25 |  |
| Medium | 24(34%) | 2.39±1.03 |  | 1.77±1.50 |  | 1.90±1.08 |  | 2.02±1.21 |  | 1.12±0.86 |  | 1.15±0.91 |  |
| High | 25(36%) | 1.99±1.39 |  | 1.81±1.44 |  | 2.06±1.19 |  | 2.36±1.38 |  | 1.32±1.11 |  | 1.19±1.03 |  |
| Smoked food: |  |  | 0.494 |  | 0.697 |  | 0.013* |  | 0.146 |  | 0.017* |  | 0.011* |
| Consumers | 46(66%) | 2.29±1.27 |  | 1.69±1.46 |  | 2.18±1.12 |  | 2.04±1.34 |  | 1.16±0.96 |  | 1.14±1.05 |  |
| Nonconsumers | 24(34%) | 2.07±1.34 |  | 1.48±1.30 |  | 1.50±0.89 |  | 1.57±1.17 |  | 1.64±0.89 |  | 1.77±1.05 |  |
| Khar: |  |  | 0.883 |  | 0.965 |  | 0.427 |  | 0.413 |  | 0.109 |  | 0.071 |
| Consumers | 47(67%) | 2.23±1.19 |  | 1.56±1.32 |  | 1.87±1.11 |  | 1.81±1.35 |  | 1.22±0.97 |  | 1.20±1.07 |  |
| Nonconsumers | 23(33%) | 2.18±1.50 |  | 1.73±1.58 |  | 2.10±1.05 |  | 2.02±1.18 |  | 1.55±0.92 |  | 1.68±1.05 |  |
| Amount of khar: |  |  | 0.687 |  | 0.850 |  | 0.696 |  | 0.380 |  | 0.121 |  | 0.072 |
| No | 23(33%) | 2.18±1.50 |  | 1.73±1.58 |  | 2.10±1.05 |  | 2.02±1.18 |  | 1.55±0.92 |  | 1.68±1.05 |  |
| Low | 23(33%) | 2.47±1.13 |  | 1.61±1.28 |  | 1.74±1.21 |  | 1.63±1.22 |  | 1.31±0.75 |  | 1.06±0.82 |  |
| Medium | 09(13%) | 1.99±1.39 |  | 1.84±1.66 |  | 1.90±1.15 |  | 2.33±1.15 |  | 0.95±0.50 |  | 0.70±0.57 |  |
| High | 15(21%) | 2.02±1.15 |  | 1.33±1.21 |  | 2.07±0.97 |  | 1.78±1.62 |  | 1.25±1.41 |  | 1.71±1.45 |  |

Note: Data are represented as mean ± standard deviation. \* Statistically significant p value.

**Table 3.** Tabulation of association of IL8, IL12 and IL13 expression with different clinicopathological parameters and lifestyle factors in female ESCC patients.

| Clinicopathological parameters and lifestyle factors | No of cases (%) | IL8 tissue |  | IL8 blood |  | IL13 tissue |  | IL13 blood |  | IL12 tissue |  | IL12 blood |  |
| --- | --- | --- | --- | --- | --- | --- | --- | --- | --- | --- | --- | --- | --- |
|  |  | mRNA expression | P value | mRNA expression | P value | mRNA expression | P value | mRNA expression | P value | mRNA expression | P value | mRNA expression | P value |
| Age group: |  |  | 0.002* |  | 0.035* |  | 0.394 |  | 0.777 |  | 0.009* |  | 0.037* |
| <50years | 18(36%) | 1.61±1.21 |  | 1.03±1.26 |  | 1.81±1.22 |  | 1.34±1.03 |  | 1.56±0.74 |  | 1.76±1.03 |  |
| >50years | 32(64%) | 2.73±1.15 |  | 1.77±1.38 |  | 2.09±0.97 |  | 1.55±1.21 |  | 0.97±0.73 |  | 1.19±0.86 |  |
| Location of tumor: |  |  | 0.655 |  | 0.612 |  | 0.261 |  | 0.133 |  | 0.948 |  | 0.217 |
| Upper esophagus | 19(38%) | 2.43±1.30 |  | 1.31±1.23 |  | 1.72±0.98 |  | 1.17±0.98 |  | 1.16±0.81 |  | 1.44±1.06 |  |
| Middle esophagus | 21(42%) | 2.13±1.25 |  | 1.39±1.17 |  | 2.04±1.03 |  | 1.55±1.28 |  | 1.23±0.74 |  | 1.53±0.76 |  |
| Lower esophagus | 10(20%) | 2.54±1.38 |  | 2.11±1.92 |  | 2.40±1.24 |  | 1.89±1.06 |  | 1.14±0.87 |  | 1.01±1.11 |  |
| Histopathology grade: |  |  | 0.810 |  | 0.591 |  | 0.019* |  | 0.005* |  | 0.559 |  | 0.858 |
| Grade 1:well differentiated ESCC | 12(24%) | 2.12±1.13 |  | 1.37±1.24 |  | 1.65±0.79 |  | 0.84±0.52 |  | 1.35±0.86 |  | 1.52±1.12 |  |
| Grade 2: moderately differentiated ESCC | 30(60%) | 2.37±1.40 |  | 1.70±1.50 |  | 1.87±1.06 |  | 1.35±0.88 |  | 1.18±0.83 |  | 1.37±0.93 |  |
| Grade 3:poorly differentiated ESCC | 08(16%) | 2.48±1.14 |  | 0.96±0.97 |  | 2.92±1.03 |  | 2.89±1.57 |  | 0.95±0.35 |  | 1.29±0.90 |  |
| Dysphagia grade: |  |  | 0.020* |  | 0.045* |  | 0.074 |  | 0.438 |  | 0.981 |  | 0.052 |
| Grade 0: Asymptomatic | 02(04%) | 0.82±0.26 |  | 0.56±0.69 |  | 1.11±0.56 |  | 3.32±2.20 |  | 1.00±0.04 |  | 2.45±0.01 |  |
| Grade 1: Solids with some dysphagia | 15(30%) | 1.87±1.19 |  | 0.83±0.79 |  | 1.93±0.80 |  | 1.48±1.23 |  | 1.27±0.92 |  | 1.75±0.92 |  |
| Grade 2: Soft, pureed food only | 11(22%) | 2.24±1.24 |  | 1.18±0.78 |  | 1.82±1.18 |  | 1.25±1.04 |  | 1.12±0.72 |  | 1.50±1.16 |  |
| Grade 3:Liquids only | 15(30%) | 2.49±1.28 |  | 2.04±1.60 |  | 1.82±1.21 |  | 1.37±0.77 |  | 1.20±0.77 |  | 1.13±0.75 |  |
| Grade 4: No<br>swallowing at all | 07(14%) | 3.52±0.78 |  | 2.55±1.81 |  | 3.00±0.62 |  | 1.54±1.36 |  | 1.11±0.82 |  | 0.72±0.72 |  |
| Tumor stage: |  |  | 0.001* |  | 0.005* |  | 0.003* |  | 0.788 |  | 0.128 |  | 0.035* |
| Stage 1 | 02(04%) | 0.99±1.05 |  | 0.13±0.06 |  | 0.68±0.19 |  | 1.41±0.08 |  | 1.90±0.03 |  | 2.83±0.64 |  |
| Stage 2 | 13(26%) | 1.73±1.17 |  | 0.73±0.66 |  | 1.60±0.86 |  | 1.79±1.54 |  | 1.25±0.71 |  | 1.72±0.91 |  |
| Stage 3 | 27(54%) | 2.29±1.19 |  | 1.57±1.23 |  | 1.96±1.05 |  | 1.29±0.91 |  | 1.26±0.85 |  | 1.30±0.93 |  |
| Stage 4 | 08(16%) | 3.76±0.48 |  | 2.87±1.75 |  | 3.05±0.71 |  | 1.62±1.28 |  | 0.66±0.43 |  | 0.80±0.70 |  |
| Node stage: |  |  | 0.574 |  | 0.312 |  | 0.258 |  | 0.646 |  | 0.136 |  | 0.814 |
| Stage 0 | 11(22%) | 2.03±1.36 |  | 1.67±1.24 |  | 1.65±0.91 |  | 1.02±0.52 |  | 1.61±0.82 |  | 1.60±1.12 |  |
| Stage 1 | 22(44%) | 2.20±1.33 |  | 1.06±1.10 |  | 1.89±1.05 |  | 0.57±1.02 |  | 0.94±0.78 |  | 1.41±0.92 |  |
| Stage 2 | 12(22%) | 2.67±1.00 |  | 1.84±1.71 |  | 2.13±1.17 |  | 1.85±1/73 |  | 1.28±0.59 |  | 1.32±0.98 |  |
| Stage 3 | 06(12%) | 2.70±1.51 |  | 2.20±1.65 |  | 2.69±1.06 |  | 1.29±1.02 |  | 1.13±0.81 |  | 1.08±0.84 |  |
| Metastasis: |  |  | 0.298 |  | 0.066 |  | 0.213 |  | 0.191 |  | 0.987 |  | 0.035* |
| Absent | 36(72%) | 2.21±1.25 |  | 1.24±1.23 |  | 1.87±1.04 |  | 1.41±1.21 |  | 1.19±0.77 |  | 1.58±0.96 |  |
| Present | 14(28%) | 2.63±1.35 |  | 2.16±1.53 |  | 2.29±1.09 |  | 1.64±0.95 |  | 1.18±0.83 |  | 0.91±0.78 |  |
| Smoking: |  |  | 0.495 |  | 0.733 |  | 0.015* |  | 0.570 |  | 0.203 |  | 0.283 |
| Smokers | 09(18%) | 2.60±1.40 |  | 1.58±1.44 |  | 2.76±0.70 |  | 1.57±1.14 |  | 0.88±0.98 |  | 1.03±0.93 |  |
| Nonsmokers | 41(82%) | 2.27±1.26 |  | 1.48±1.38 |  | 1.82±1.06 |  | 1.46±1.15 |  | 1.25±0.73 |  | 1.47±0.95 |  |
| Alcohol: |  |  | 0.161 |  | 0.686 |  | 0.063 |  | 0.771 |  | 0.159 |  | 0.438 |
| Alcoholic | 05(10%) | 3.09±1.36 |  | 1.79±1.81 |  | 2.83±0.84 |  | 1.31±0.59 |  | 0.71±0.53 |  | 1.01±0.96 |  |
| Nonalcoholic | 45(90%) | 2.24±1.26 |  | 1.47±1.34 |  | 1.89±1.05 |  | 1.50±1.19 |  | 1.24±0.79 |  | 1.44±0.96 |  |
| Betel nut: |  |  | 0.041* |  | 0.023* |  | 0.879 |  | 0.199 |  | 0.014* |  | 0.604 |
| Chewers | 36(72%) | 2.56±1.30 |  | 1.77±1.45 |  | 1.97±1.14 |  | 1.60±1.21 |  | 1.02±0.73 |  | 1.34±0.94 |  |
| Nonchewers | 14(28%) | 1.73±1.06 |  | 0.80±0.83 |  | 2.03±0.86 |  | 1.15±0.88 |  | 1.62±0.77 |  | 1.52±1.03 |  |
| Tobacco: |  |  | 0.690 |  | 0.821 |  | 0.024* |  | 0.017* |  | 0.082 |  | 0.284 |
| Chewers | 29(58%) | 2.39±1.24 |  | 1.44±1.29 |  | 2.27±1.14 |  | 1.83±1.28 |  | 1.02±0.82 |  | 1.26±0.93 |  |
| Nonchewers | 21(42%) | 2.24±1.37 |  | 1.58±1.50 |  | 1.59±0.82 |  | 0.99±0.68 |  | 1.41±0.67 |  | 1.58±0.96 |  |
| Fish: |  |  | 0.754 |  | 0.482 |  | 0.022* |  | 0.065 |  | 0.266 |  | 0.438 |
| Consumers | 40(80%) | 2.36±1.28 |  | 1.56±1.36 |  | 2.16±1.09 |  | 1.62±1.21 |  | 1.12±0.79 |  | 1.34±0.92 |  |
| Nonconsumers | 10(20%) | 2.21±1.37 |  | 1.26±1.46 |  | 1.30±0.60 |  | 0.89±0.53 |  | 1.43±0.74 |  | 1.62±1.09 |  |
| Meat: |  |  | 0.824 |  | 0.615 |  | 0.035* |  | 0.078 |  | 0.247 |  | 0.128 |
| Consumers | 43(86%) | 2.34±1.28 |  | 1.46±1.36 |  | 2.12±1.09 |  | 1.59±1.17 |  | 1.37±0.77 |  | 1.31±0.95 |  |
| Nonconsumers | 07(14%) | 2.22±1.40 |  | 1.73±1.56 |  | 1.20±0.37 |  | 0.79±0.60 |  | 1.51±0.82 |  | 1.88±0.94 |  |
| Egg: |  |  | 0.321 |  | 0.884 |  | 0.439 |  | 0.961 |  | 0.800 |  | 0.910 |
| Consumers | 45(90%) | 2.27±1.29 |  | 1.48±1.32 |  | 2.03±1.06 |  | 1.48±1.14 |  | 1.19±0.80 |  | 1.39±0.98 |  |
| Nonconsumers | 05(10%) | 2.87±1.17 |  | 1.70±2.00 |  | 1.63±1.19 |  | 1.48±1.22 |  | 1.10±0.59 |  | 1.44±0.76 |  |
| Pickle: |  |  | 0.804 |  | 0.382 |  | 0.856 |  | 0.578 |  | 0.628 |  | 0.508 |
| Consumers | 42(84%) | 2.35±1.38 |  | 1.59±1.43 |  | 1.98±1.10 |  | 1.45±1.16 |  | 1.21±0.79 |  | 1.43±0.98 |  |
| Nonconsumers | 08(16%) | 2.22±0.55 |  | 1.03±0.94 |  | 2.05±0.88 |  | 1.62±1.11 |  | 1.06±0.73 |  | 1.18±0.84 |  |
| Spices: |  |  | 0.031* |  | 0.041* |  | 0.962 |  | 0.261 |  | 0.004* |  | 0.034* |
| Consumers | 36(72%) | 2.57±1.27 |  | 1.76±1.48 |  | 1.98±1.10 |  | 1.62±1.24 |  | 0.99±0.65 |  | 1.20±0.93 |  |
| Nonconsumers | 14(28%) | 1.70±1.13 |  | 0.83±0.75 |  | 2.00±1.00 |  | 1.12±0.73 |  | 1.68±0.89 |  | 1.88±0.87 |  |
| Fast food: |  |  | 0.976 |  | 0.674 |  | 0.374 |  | 0.324 |  | 0.427 |  | 0.860 |
| Consumers | 22(44%) | 2.32±1.33 |  | 1.63±1.49 |  | 2.14±0.98 |  | 1.67±1.23 |  | 1.08±0.70 |  | 1.41±0.95 |  |
| Nonconsumers | 28(56%) | 2.33±1.27 |  | 1.40±1.29 |  | 1.79±1.14 |  | 1.33±1.06 |  | 1.26±0.84 |  | 1.38±0.98 |  |
| Hot food: |  |  | 0.022* |  | 0.033* |  | 0.203 |  | 0.547 |  | 0.034* |  | 0.062 |
| Consumers | 25(50%) | 2.74±1.26 |  | 1.97±1.49 |  | 2.18±0.97 |  | 1.36±1.01 |  | 0.95±0.73 |  | 1.14±0.88 |  |
| Nonconsumers | 25(50%) | 1.92±1.19 |  | 1.04±1.08 |  | 1.79±1.14 |  | 1.60±1.27 |  | 1.42±0.77 |  | 1.65±0.98 |  |
| Type of tea: |  |  | 0.189 |  | 0.595 |  | 0.149 |  | 0.512 |  | 0.133 |  | 0.175 |
| Red tea | 08(16%) | 2.43±1.02 |  | 1.08±1.50 |  | 1.65±0.97 |  | 1.21±1.03 |  | 1.61±0.69 |  | 1.37±1.05 |  |
| Milk tea | 21(42%) | 1.95±1.51 |  | 1.56±1.29 |  | 1.77±1.04 |  | 1.80±1.43 |  | 1.24±0.88 |  | 1.70±0.99 |  |
| Both | 21(42%) | 2.67±1.04 |  | 1.60±1.44 |  | 2.33±1.07 |  | 1.26±0.75 |  | 0.97±0.65 |  | 1.10±0.83 |  |
| Amount of tea: |  |  | 0.011* |  | 0.594 |  | 0.248 |  | 0.689 |  | 0.647 |  | 0.047* |
| Low | 20(40%) | 2.63±1.21 |  | 1.62±1.52 |  | 1.95±1.05 |  | 1.46±1.08 |  | 1.26±0.92 |  | 1.51±0.92 |  |
| Medium | 15(30%) | 1.52±1.14 |  | 1.23±1.10 |  | 1.68±0.78 |  | 1.35±1.20 |  | 1.24±0.67 |  | 1.75±1.04 |  |
| High | 15(30%) | 2.73±1.21 |  | 1.61±1.46 |  | 2.34±1.27 |  | 1.63±1.22 |  | 1.03±0.70 |  | 0.89±0.73 |  |
| Smoked food: |  |  | 0.042* |  | 0.938 |  | 0.037* |  | 0.837 |  | 0.022* |  | 0.029* |
| Consumers | 28(56%) | 2.65±1.35 |  | 1.57±1.46 |  | 2.27±1.07 |  | 1.53±1.28 |  | 0.96±0.72 |  | 1.12±0.86 |  |
| Nonconsumers | 22(44%) | 1.91±1.08 |  | 1.41±1.29 |  | 1.63±0.97 |  | 1.41±0.96 |  | 1.47±0.77 |  | 1.75±0.97 |  |
| Khar: |  |  | 0.282 |  | 0.409 |  | 0.593 |  | 0.019* |  | 0.360 |  | 0.081 |
| Consumers | 35(70%) | 2.46±1.33 |  | 1.60±1.41 |  | 1.93±1.11 |  | 1.73±1.25 |  | 1.12±0.80 |  | 1.26±0.97 |  |
| Nonconsumers | 15(30%) | 2.02±1.13 |  | 1.28±1.32 |  | 2.11±0.97 |  | 0.89±0.45 |  | 1.34±0.73 |  | 1.72±0.86 |  |
| Amount of khar: |  |  | 0.655 |  | 0.721 |  | 0.219 |  | 0.007* |  | 0.641 |  | 0.148 |
| No | 15(30%) | 2.02±1.13 |  | 1.28±1.32 |  | 2.11±0.97 |  | 0.89±0.45 |  | 1.34±0.73 |  | 1.72±0.86 |  |
| Low | 17(34%) | 2.46±1.41 |  | 1.59±1.45 |  | 1.74±0.98 |  | 1.22±0.72 |  | 1.13±0.84 |  | 1.50±1.10 |  |
| Medium | 09(18%) | 2.66±1.32 |  | 2.01±1.59 |  | 2.21±0.96 |  | 1.47±1.04 |  | 0.93±0.79 |  | 0.99±0.87 |  |
| High | 09(18%) | 2.24±1.32 |  | 1.19±1.14 |  | 2.03±1.49 |  | 2.94±1.53 |  | 1.28±0.78 |  | 1.05±0.78 |  |

Note: Data are represented as mean ± standard deviation. \* Statistically significant p value.

### Relationship between IL8, IL12 and IL13 expression in tissue and blood samples of ESCC patients

Significant positive correlation (p<0.05) was observed between tissue and blood level expression of IL8, IL12 and IL13. Again, a significant positive correlation of IL8 and IL13 expression and a significant negative correlation of IL8 and IL12 expression were revealed when correlation was performed between the two cytokines in both tissue and blood level. Correlation study of IL8, IL12 and IL13 gene expression in tissue and blood level are represented in Table 4 and S13 Fig.

**Table 4.** Correlation study of IL8, IL12 and IL13 gene expression in tissue and blood level.

| Parameters | P value | Correlation |
| --- | --- | --- |
| IL8 tissue and IL8 blood | 0.000* | 0.492 |
| IL13 tissue and IL13 blood | 0.017* | 0.217 |
| IL12 tissue and IL12 blood | 0.004* | 0.262 |
| IL8 tissue and IL13 tissue | 0.047* | 0.182 |
| IL8 tissue and IL12 tissue | 0.000* | -0.372 |
| IL13 tissue and IL12 tissue | 0.082 | -0.159 |
| IL8 blood and IL13 blood | 0.001* | 0.301 |
| IL8 blood and IL12 blood | 0.023* | -0.208 |
| IL13 blood and IL12 blood | 0.651 | -0.042 |

Note: \* Statistically significant p value.

### Survival analysis

Among males, tumor stage 4 was observed with lower mean survival time than other tumor stages (p=0.012). Similarly, males possessing node stage 3 were noted with lower survival time than other node stages (p=0.009). Metastatic male patients were observed to have a lower survival than nonmetastatic males (p=0.024). Males consuming alcohol, hot food and smoked food in their diet were noticed with lower survival time than the others who don’t consume the same (p=0.021, p=0.013 and p=0.021 respectively). Again, males having lower levels of IL12 in their tissue were observed to have lower survival compared to males with higher IL12 expression (p=0.005).

Females possessing histology grade 3 were noticed to have lower survival compared to histopathology grade 1 and 2 (p=0.024). Similarly, females possessing tumor stage 1 were noted with higher mean survival time than other tumor stages (p=0.015). Metastatic females were observed with a lower survival time than non-metastatic one (p=0.022). Females consuming betel nut, tobacco, spices, hot food and smoked food were noticed with lower survival time than the others who don’t consume the same (p=0.030, p=0.015, p=0.015, p=0.004 and p=0.015 respectively). Females having lower levels of IL12 in their tissue and blood were observed to have lower survival compared to females with higher IL12 expression in their tissue and blood (p=0.003 and p=0.004 respectively). Similarly, females having higher levels of IL8 expression in their blood and higher levels of IL13 expression in their tissue were noted to have lower survival (p=0.016 and p=0.031 respectively). Survival analysis data for males and females are listed in Table 5.

**Table 5.**
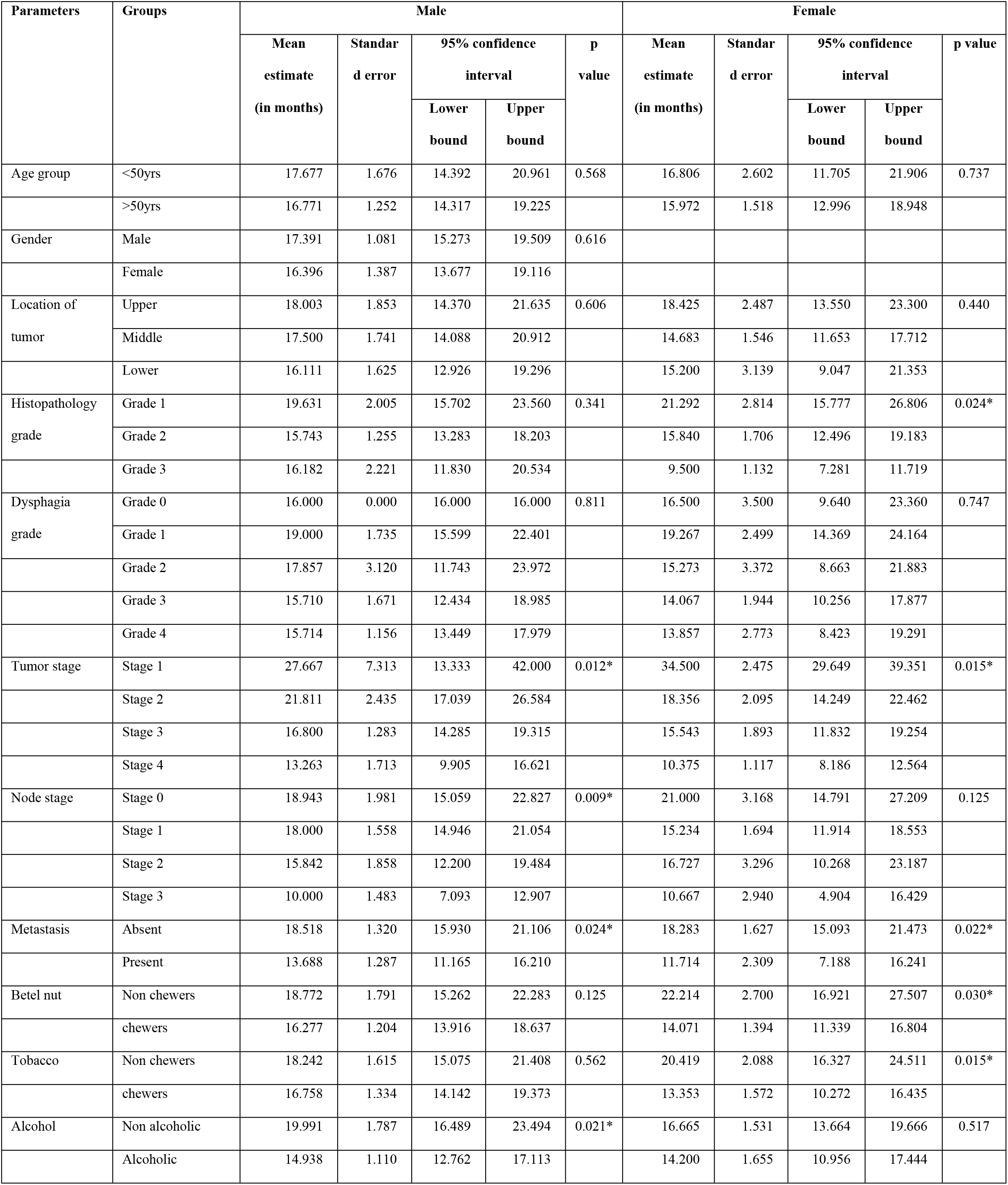
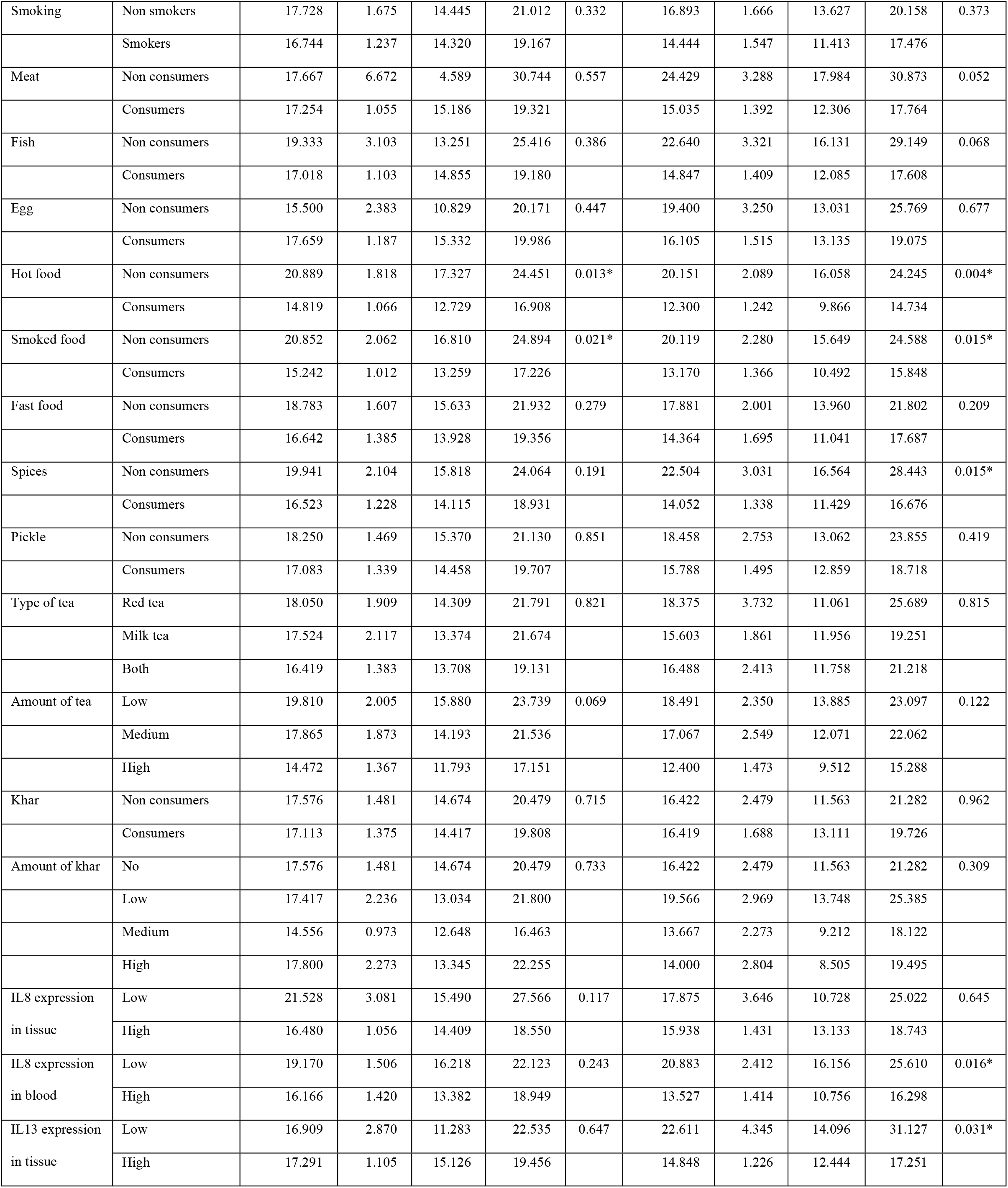
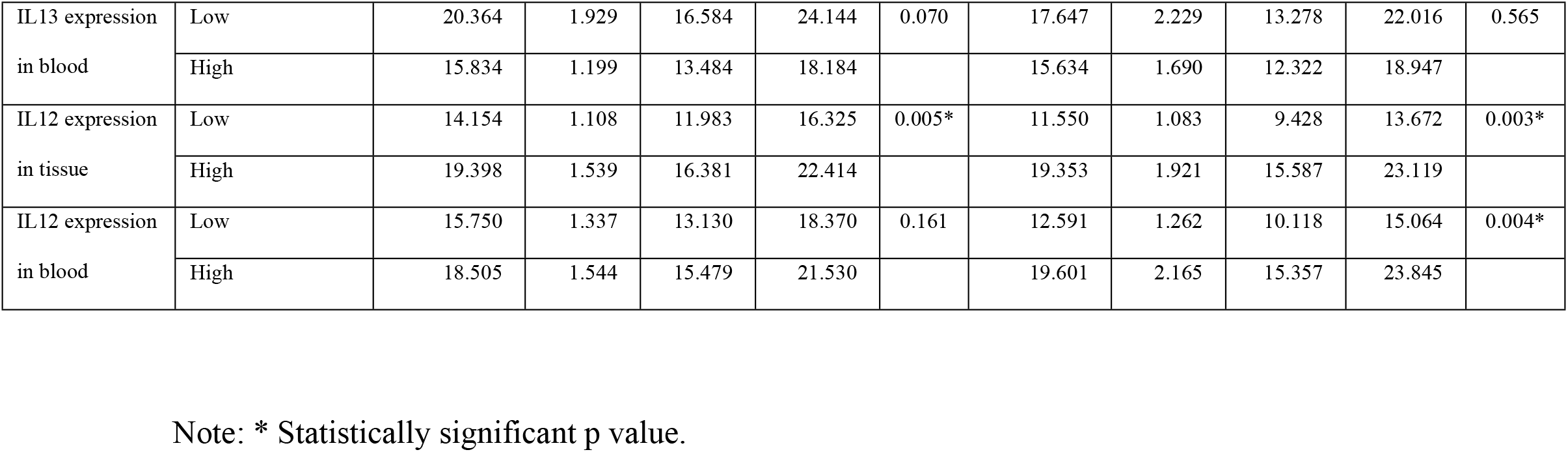
Tabulation of survival analysis data of both male and female ESCC patients.

### Hazard analysis

The hazard (mortality) ratio is 4.649 times higher for males having node stage 3 compared to node stage 0 (p=0.004). The hazard ratio is higher for males having metastasis compared to nonmetastatic one (p=0.036). Similarly, the hazard ratio is higher for males consuming alcohol, hot food and smoked food in their diet compared to one who don’t consume the same (p=0.032, p=0.021 and p=0.032 respectively). The hazard ratio is higher for males consuming higher amount of tea compared to one consuming lower amount of tea (p=0.037). Moreover, the hazard ratio is 0.499 times for male patients with higher levels of IL12 in their tissue compared to one with a lower level of IL12 expression (p=0.009).

Among females, significant difference in hazard ratio was observed between histopathology grade 1 and histopathology grade 3 (p=0.012). The hazard ratio is higher for females having tumor stage 4 compared to tumor stage 1 (p=0.014). Similarly, the hazard ratio is higher for females having node stage 3 compared to node stage 0 (p=0.029). The hazard ratio for a metastatic female patient is 2.064 times that of a nonmetaststic one (P=0.032). The hazard ratio is significantly higher in females consuming betel nut, tobacco, hot food, smoked food and spices compared to others who don’t consume the same (p=0.042, p=0.023, p=0.007, p=0.023 and p=0.023 respectively). Moreover, the hazard ratio is higher for female patients with higher levels of IL8 expression in their blood samples and higher levels of IL13 expression in their tissue samples (p=0.023 and p=0.045 respectively). Again, the hazard ratio is 0.404 times and 0.425 times for female patients with higher levels of IL12 in their tissue and blood compared to one with a lower level of IL12 expression (p=0.007 and p=0.008 respectively). Hazard analysis data for males and females are listed in Table 6. Survival and hazard analysis graphs representing different parameters are presented in S14-S20 Figs for male patients and S21-S28 Figs for female patients.

**Table 6.**
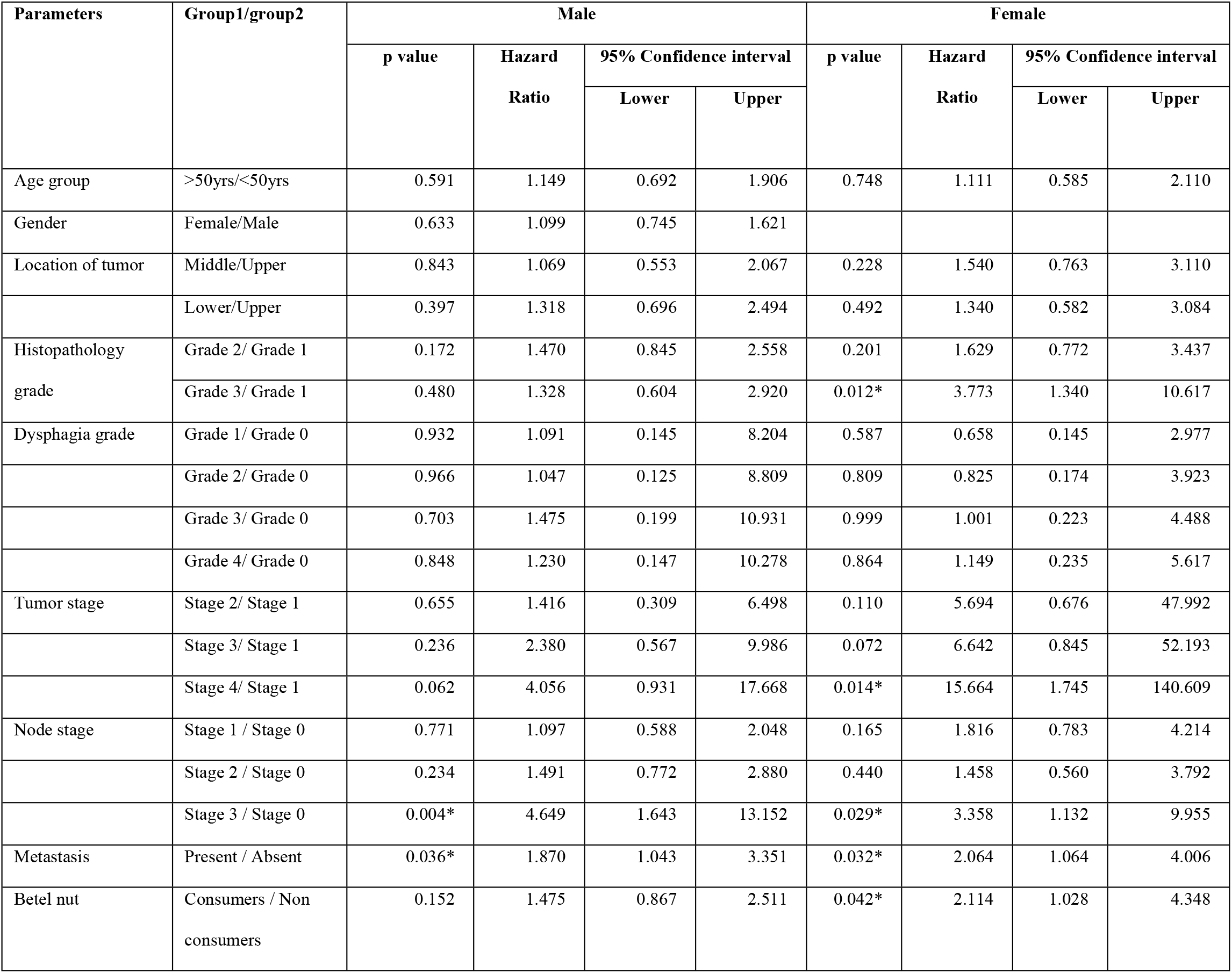
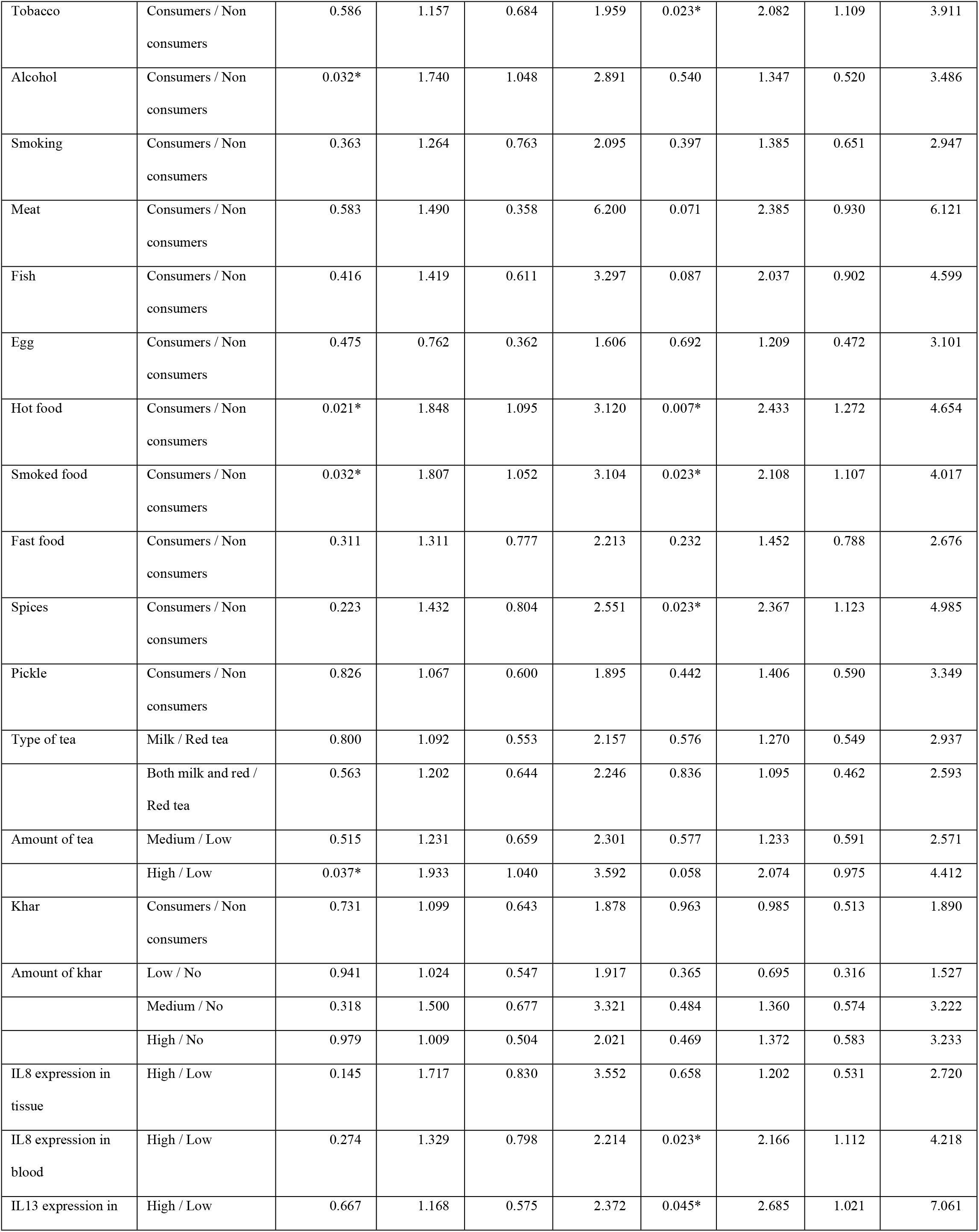
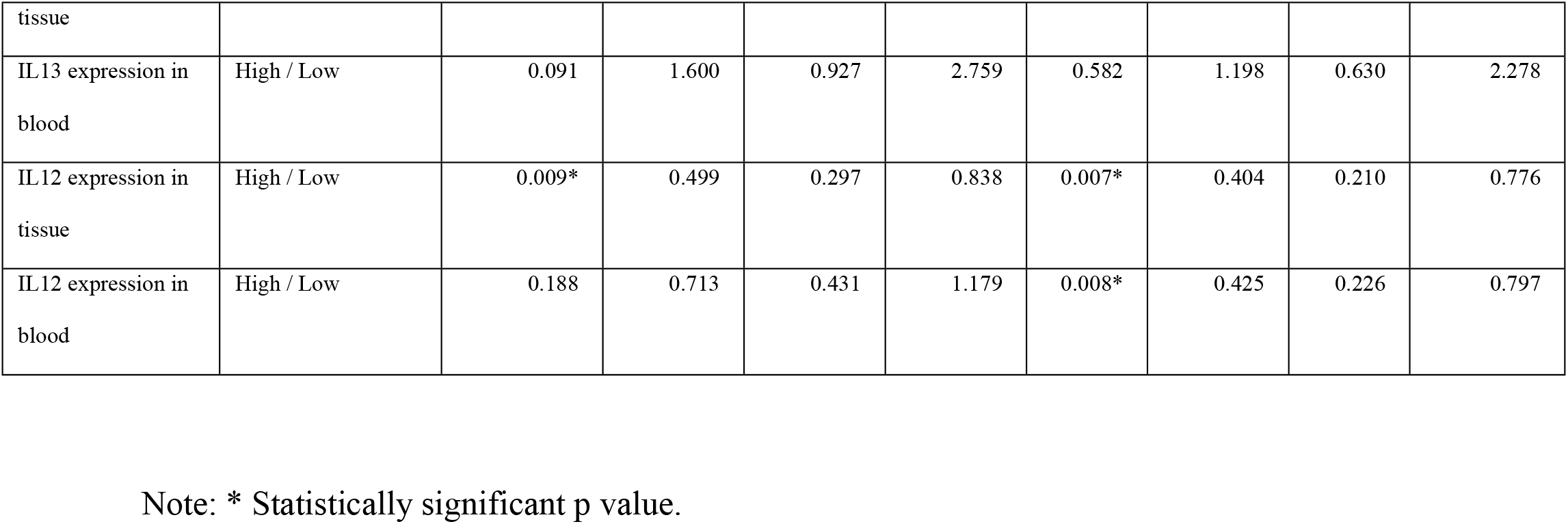
Tabulation of hazard analysis data of both male and female ESCC patients.

## Discussion

This study targeted both tissue and blood samples of the targeted patients for analysis of expression profile of the selected cytokines. This study of gene expression at the blood and tissue levels was done independently for men and women. Till date very less expression studies were performed together in both tissue and blood samples.

High level of IL8 expression was observed in many human cancers, including breast, lung, prostate, pancreatic, colorectal, esophageal cancers as well as melanoma and numerous studies have shown that serum level of IL-8 in cancer patients can act like a prognostic marker [21-25]. Increased IL8 expression was observed in gastric cancer tissue and overexpression of IL-8 was reported to have association with prognosis of this cancer [26]. Higher level of IL8 was also reported in the sera of liver, gastric and non-small-cell lung cancer patients as compared to healthy controls [27]. Moreover, overexpression of IL8 was correlated with tumor progression, recurrence and the TNM stage in multiple cancers [28]. In our study, 83% patients showed overexpression of IL8 in their tissue (p=0.000). This data reveal higher potential of IL8 expression in tissue samples as prospective molecular biomarker for screening ESCC. But only 62% patients were noted to have higher levels of IL8 expression in their blood (p=0.388). Moreover, a significant positive correlation (p=0.000) between blood and tissue level IL8 expression was also observed in this study.

Higher level of IL12 was reported in the sera of non-small-cell lung cancer, prostate carcinoma and metastatic renal cell carcinoma patients as compared to healthy controls [27, 29]. But lower level of IL12 was reported in the serum of breast cancer patients when compared to healthy controls [30]. Higher level of IL12 was also reported in the serum of esophageal SCC patients as compared to controls [31]. In our study, high level of IL12 expression was observed in 57% blood samples (p=0.222) and 62% tissue samples (p=0.435). A significant positive correlation (p=0.004) between blood and tissue level IL12 expression was also observed in ESCC patients.

Increased expression of IL13 was reported in peripheral blood of breast, prostate and bladder cancer patients [32]. High level of IL13 expression was noticed in pancreatic cancer, lymphoma, oral squamous cell carcinoma and non-small cell lung Carcinoma patients [33-36]. Plasma level of IL-13 was significantly higher in bladder cancer patients than in the healthy controls and higher serum IL-13 level was reported to have association with progression of diffuse large B cell lymphoma [37, 38]. Higher concentration of IL-13 was observed in colorectal and upper gastrointestinal tract tumors than adjacent normal tissue [39]. Higher level of IL13 was also reported in the sera of melanoma and skin cancer patients as compared to healthy controls [27]. In our study, high level of IL13 expression was observed in 68% blood samples (p=0.312) and 83% tissue samples (p=0.001). These data indicate a higher potential of IL13 expression in tissue samples as prospective molecular biomarker for screening ESCC. Additionally, a significant positive correlation (p=0.017) between blood and tissue level IL13 expression was also observed in this study.

In India, the role of diet, nutrition and food habits in causing esophageal cancer has been given attention recently. Consumption of very hot foods, spices, smoked food and some locally made food e.g. khar (kalakhar), a locally made food of Assam, were reported to have significant associations with the risk esophageal cancer development [40-42]. Betel nut chewing, consumption of tobacco (smoking and smokeless tobacco) and alcohol were noted to have significant association with an increased risk of esophageal cancer [40, 42, 43]. In our study, different clinicopathological factors and dietary habits like age group, dysphagia grade, histopathology grade, consumption of betel nut, tobacco, hot food, smoked food, spices, etc. showed significant association (p<0.05) with IL8, IL13 and IL12 expression in tissue and blood samples of ESCC patients. These data represent their association and clinical significance with the alteration of the studied cytokines’ expression in ESCC.

Although esophageal cancer prevention approaches are essential, measures to lower morbidity or increase survival are equally crucial. Different clinicopathological parameters and risk factors like histopathology grade, age, gender, tumor stage etc. affect the survival of ESCC patients [20, 44, 45]. In our study, tumor stage, node stage, metastasis, consumption of hot food, smoked food and alcohol showed significant association (p<0.05) with survival of male patients, whereas survival of female patients showed association (p<0.05) with histopathology grade, tumor stage, metastasis, consumption of betel nut, spices, smoked food, hot food and tobacco. Moreover, the univariate model of hazard analysis data also supports these findings and all these data represent their clinical importance for detecting survival of ESCC patients. When the expression of IL8, IL13, and IL12 was analysed with survival and hazard outcomes, statistically significant association was observed in IL8 expression in female patient blood, IL13 expression in female patient tissue, and IL12 expression in male and female patient tissue as well as in female patient blood. This clearly illustrates the function of these cytokines and their clinical significance in determining ESCC patient survival.

The findings of our study show some similarities and differences with those of other studies, which may be caused by variations in sample size, geographic location, genetic and environmental factors, racial and ethnic diversity, associated clinical conditions, etc. The distinct genetic makeup and indigenous food habit among the Northeast Indian population may be factors for higher incidence of esophageal cancer in this area. Different risk factors or variables like individual diet, nutrition, lifestyle, food habits, genetic, epigenetic and environmental factors, etc. determine which type of cancer predominates in a given patient or in a given geographical location [40, 42, 46]. This study will provide us to acquire more knowledge towards the role of IL8, IL12 and IL13 on ESCC progression along with its interaction with different clinicopathological factors and dietary habits for causing this cancer in the Northeast Indian population.

In conclusion, altered expression of IL8, IL12 and IL13 may be associated with ESCC progression. This expression study also reveals the correlation of studied cytokines in tissue and blood level and the association of different clinicopathological and dietary factors in ESCC. Moreover, both survival and hazard analysis data also reveals the impact of different factors on survival and mortality rate of ESCC patients. Again, overexpression of both IL8 and IL13 in tissue samples may be a potential biomarker for ECSS screening among Northeast Indian Population. This type of gene expression study along with survival and hazard outcomes will enable us to learn more about and develop a deeper understanding of the biology of esophageal cancer.

## Supporting information

**S1 Fig. Box plot representation of significant association (p<0.05) of IL8 expression in tissue level with different parameters in ESCC male patients**.

**S2 Fig. Box plot representation of significant association (p<0.05) of IL8 expression in tissue level with different parameters in ESCC female patients**.

**S3 Fig. Box plot representation of significant association (p<0.05) of IL8 expression in blood level with different parameters in ESCC male patients**.

**S4 Fig. Box plot representation of significant association (p<0.05) of IL8 expression in blood level with different parameters in ESCC female patients**.

**S5 Fig. Box plot representation of significant association (p<0.05) of IL12 expression in tissue level with different parameters in ESCC male patients**.

**S6 Fig. Box plot representation of significant association (p<0.05) of IL12 expression in tissue level with different parameters in ESCC female patients**.

**S7 Fig. Box plot representation of significant association (p<0.05) of IL12 expression in blood level with different parameters in ESCC male patients**.

**S8 Fig. Box plot representation of significant association (p<0.05) of IL12 expression in blood level with different parameters in ESCC female patients**.

**S9 Fig. Box plot representation of significant association (p<0.05) of IL13 expression in tissue level with different parameters in ESCC male patients**.

**S10 Fig. Box plot representation of significant association (p<0.05) of IL13 expression in tissue level with different parameters in ESCC female patients**.

**S11 Fig. Box plot representation of significant association (p<0.05) of IL13 expression in blood level with different parameters in ESCC male patients**.

**S12 Fig. Box plot representation of significant association (p<0.05) of IL13 expression in blood level with different parameters in ESCC female patients**.

**S13 Fig. Scattered plot representation of correlation of IL8, IL12 and IL13 expression in tissue and blood level**

**S14 Fig. Graphical representation of survival and hazard analysis in males representing different parameters**.

**S15 Fig. Graphical representation of survival and hazard analysis in males representing IL8 expression in tissue level**.

**S16 Fig. Graphical representation of survival and hazard analysis in males representing IL8 expression in blood level**.

**S17 Fig. Graphical representation of survival and hazard analysis in males representing IL12 expression in tissue level**.

**S18 Fig. Graphical representation of survival and hazard analysis in males representing IL12 expression in blood level**.

**S19 Fig. Graphical representation of survival and hazard analysis in males representing IL13 expression in tissue level**.

**S20 Fig. Graphical representation of survival and hazard analysis in males representing IL13 expression in blood level**.

**S21 Fig. Graphical representation of survival and hazard analysis in females representing different parameters**.

**S22 Fig. Graphical representation of survival and hazard analysis in ESCC patients representing gender**.

**S23 Fig. Graphical representation of survival and hazard analysis in females representing IL8 expression in tissue level**.

**S24 Fig. Graphical representation of survival and hazard analysis in females representing IL8 expression in blood level**.

**S25 Fig. Graphical representation of survival and hazard analysis in females representing IL13 expression in tissue level**.

**S26 Fig. Graphical representation of survival and hazard analysis in females representing IL13 expression in blood level**.

**S27 Fig. Graphical representation of survival and hazard analysis in females representing IL12 expression in tissue level**.

**S28 Fig. Graphical representation of survival and hazard analysis in females representing IL12 expression in blood level**.

## Author contributions

**Conceptualization:** Jayasree Talukdar, Abdul Malik, Subhash Medhi

**Data curation**: Jayasree Talukdar, Subhash Medhi

**Funding acquisition**: Abdul Malik, Subhash Medhi

**Methodology:** Jayasree Talukdar, Abdul Malik, Manash Pratim Sarma, Minakshi Bhattacharjee, Mrinmoy Basak, Manash Pratim Kashyap, Musaed Alkholief, Aws Alshamsan, Subhash Medhi

**Project administration:** Subhash Medhi

**Software:** Jayasree Talukdar, Kangkana Kataki, Manash Pratim Kashyap, Sahana Bhattacharjee, Eyashin Ali, Chenole Keppen

**Validation:** Bikash Narayan Choudhury, Munindra Narayan Baruah, Mallika Bhattacharyya, Mrinmoy Basak, Mohammad Ghaznavi Idris, Subhash Medhi

**Formal Analysis:** Jayasree Talukdar

**Investigation:** Jayasree Talukdar, Bikash Narayan Choudhury, Munindra Narayan Baruah, Mallika Bhattacharyya, Manash Pratim Sarma, Minakshi Bhattacharjee, Mohammad Ghaznavi Idris, Subhash Medhi

**Resources:** Bikash Narayan Choudhury, Munindra Narayan Baruah, Mallika Bhattacharyya, Subhash Medhi

**Writing – original draft:** Jayasree Talukdar, Abdul Malik, Musaed Alkholief, Aws Alshamsan, Mrinmoy Basak, Manash Pratim Kashyap, Subhash Medhi

**Writing - review and editing:** Jayasree Talukdar, Abdul Malik, Musaed Alkholief, Aws Alshamsan, Kangkana Kataki, Manash Pratim Sarma, Minakshi Bhattacharjee, Subhash Medhi

**Visualization:** Jayasree Talukdar, Bikash Narayan Choudhury, Munindra Narayan Baruah, Mallika Bhattacharyya, Manash Pratim Sarma, Minakshi Bhattacharjee, Mohammad Ghaznavi Idris, Subhash Medhi

**Supervision:** Subhash Medhi

